# A novel method for measuring phenotypic colistin resistance in *Escherichia coli* populations from chicken flocks

**DOI:** 10.1101/2020.10.22.351577

**Authors:** Nhung Thi Nguyen, Nguyen Thi Phuong Yen, Nguyen Van Ky Thien, Nguyen Van Cuong, Bach Tuan Kiet, James Campbell, Guy Thwaites, Stephen Baker, Ronald B. Geskus, Juan Carrique-Mas

## Abstract

Colistin is extensively used in animal production in many low- and middle-income countries. There is a need to develop methodologies to benchmark and monitor changes in resistance in commensal bacterial populations in farms. We aimed to evaluate the performance of a broth microdilution method based on culturing a pooled *Escherichia coli* suspension (30-50 organisms) from each sample. In order to confirm the biological basis and sensitivity of the method, we prepared 16 standard suspensions containing variable ratios of colistin-susceptible and *mcr*-1 encoded colistin-resistant *E. coli* which were grown in 2mg/L colistin. The optical density (OD_600nm_) readings over time were used to generate a growth curve, and were adjusted to the values obtained in the absence of colistin. The median limit of detection of the method was 1 colistin-resistant in 10^4^ susceptible colonies [1^st^ - 3^rd^ quartile, 1:10^2^ – 1:10^5^]. We applied this method to 108 pooled faecal samples from 36 chicken flocks in the Mekong Delta (Vietnam) over the production cycle. The correlation between this method and the prevalence of colistin resistance in individual colonies harvested from field samples, determined by the Minimum Inhibitory Concentration (MIC), was established. The overall prevalence of colistin resistance at sample and isolate level was 38.9% and 19.4%, respectively. Increased colistin resistance was associated with recent (2 weeks) use of colistin and other, non-colistin antimicrobials (OR=3.67 and OR=1.84, respectively). Our method is a sensitive and affordable approach to monitor changes in colistin resistance in pooled *E. coli* populations from faecal samples over time.

**IMPORTANCE:** Colistin (polymyxin E) is an antimicrobial with poor solubility properties, and therefore broth microdilution is the only appropriate method for testing colistin resistance. However, estimating colistin resistance in commensal mixed *Escherichia coli* populations is laborious since it requires individual colony isolation, identification and susceptibility testing. We developed a growth-based microdilution method suitable for pooled faecal samples. We validated the method by comparing it with results from individual MIC testing of 909 *E. coli* isolates. We used the method to investigate phenotypic colistin resistance in 108 pooled faecal samples from 36 healthy chicken flocks, each sampled three times over the production cycle. A higher level of resistance was seen in flocks recently supplemented with colistin in drinking water, although the observed generated resistance was short-lived. Our method is affordable, and may potentially be integrated into surveillance systems aiming at estimating the prevalence of resistance at colony level in flocks/herds. Furthermore, it may also be adapted to other complex biological systems, such as farms and abattoirs.

## INTRODUCTION

Colistin (polymyxin E) is a last-resort drug used for the treatment of severe multi-drug resistant (MDR) infections in many countries, and currently is classified by the World Health Organization (WHO) as a ‘highest priority, critically important’ antimicrobial (1). The emergence of *mcr*-1 plasmid-encoded colistin resistance among Gram-negative bacteria is considered a serious threat to global health (2). It has been hypothesized that colistin use in animal production is a major contributing factor to the emergence of colistin resistance worldwide (3). Colistin is still used in poultry and pig farming in many countries (4). In terms of frequency, colistin is the most commonly used antimicrobial in chicken production in the Mekong Delta region of Vietnam (5, 6). Studies in the same region have shown that resistance against colistin in commensal *Escherichia coli* from chicken flocks is often encoded by the *mcr*-1 gene (7, 8). At sample level, the prevalence of *mcr*-1 in chicken faecal samples in the Mekong Delta was 59.4%. The prevalence of this gene has also be found to be higher among in-contact humans (chicken farmers) than in urban individuals (7).

*E. coli* is an ubiquitous commensal enteric organism globally used to monitor phenotypic antimicrobial resistance (AMR) in national surveillance programmes, both in humans and in animals (9, 10). Given the diversity of this organism within the enteric microbiome, the characterisation of phenotypic resistance in a mixed population of commensal *E. coli* requires selecting a representative and sufficiently large number of strains. This is often achieved by performing differential colony counts on agar media with and without antimicrobials (11). However, agar-based methods are not appropriate for colistin given the antimicrobials’ poor solubility (12). Determination of the minimal inhibitory concentration (MIC) by broth microdilution is regarded as the gold standard for testing of colistin resistance of Enterobacteriaceae (ISO 20776-1) both by the Clinical and Laboratory Standards Institute (CLSI) and the European Committee on Antimicrobial Susceptibility Testing (EUCAST) (12, 13). Establishing accurately the prevalence of resistance at colony level requires the investigation of a sufficiently large, representative number of isolates from each sample, which is extremely laborious and costly (8, 11, 14). Therefore, there is a need for developing cost-effective methodologies for evaluating resistance against colistin in mixed *E. coli* populations from animal faecal samples. Here, we designed and evaluated a broth microdilution-based method to quantify colistin resistance in *E. coli* populations from pooled chicken faecal samples. We then related the observed results to data on antimicrobial use (AMU) from the same flocks.

## RESULTS

### Growth of standard suspensions

The AUC_adj_ values generated from all susceptible-resistant combinations are presented in Fig.1. In all cases, AUC_adj_ values increased with increasing ratio of resistant to susceptible organisms. Growth was detected at a ratio of 1 resistant to 10^5^, 10^4^, 10^3^, 10^2^ and 10^1^ susceptible strains for 43.7%, 12.5%, 18.5% and 12.5% and 12.5% combinations, respectively. There was no difference in average AUC_adj_ between resistant strains with low (R1 and R2, MIC= 4mg/L) and moderate (R3 and R4, MIC= 8mg/L) levels of resistance (both AUC_adj_= 0.39). There were significant difference between average AUC_adj_ values using different susceptible strains, with values ranging from 0.09 to 0.62 (Kruskal Wallis test, p= 0.002). In combinations with resistant strains, S2 yielded the lowest average AUC_adj_ (median 0.09 [1^st^ - 3^rd^ quartile, 0.07-0.29]) as well as the lowest limit of detection (average S:R ratio of 10^2^:1). S4 gave the highest median AUC_adj_ (0.62 [1^st^ - 3^rd^ quartile, 0.48-0.69]) as well as the highest limit of detection (average S:R ratio of 10^5^:1).

### Study flocks and their AMU

A total of 36 flocks (108 samples) were investigated in this study. The median flock size was 231 [1^st^ - 3^rd^ quartile, 189-401] chickens. Flocks were raised over a median of 19 [1^st^ - 3^rd^ quartile, 17-20] weeks. Colistin had been administered to 22/36 (61.1%) flocks. Among flocks given colistin, the average number of Animal Daily Doses (ADD) per 1,000 chicken-days of this antimicrobial administered over the production cycle was 149.5 Standard deviation [SD] ±261.6. Colistin was used more during the early flock cycle period (281.7 SD ±321.2 ADDs per 1,000 chicken-days) compared with the second period (17.4 SD ±18.1 ADDs per 1,000 chicken-days) (Wilcoxon paired test, p<0.001) (Table 1). This antimicrobial was administrated over a median of 4 [1^st^ - 3^rd^ quartile, 2-6] weeks. The data of colistin use among study flocks is displayed in Fig. S2.

**FIG 1.**
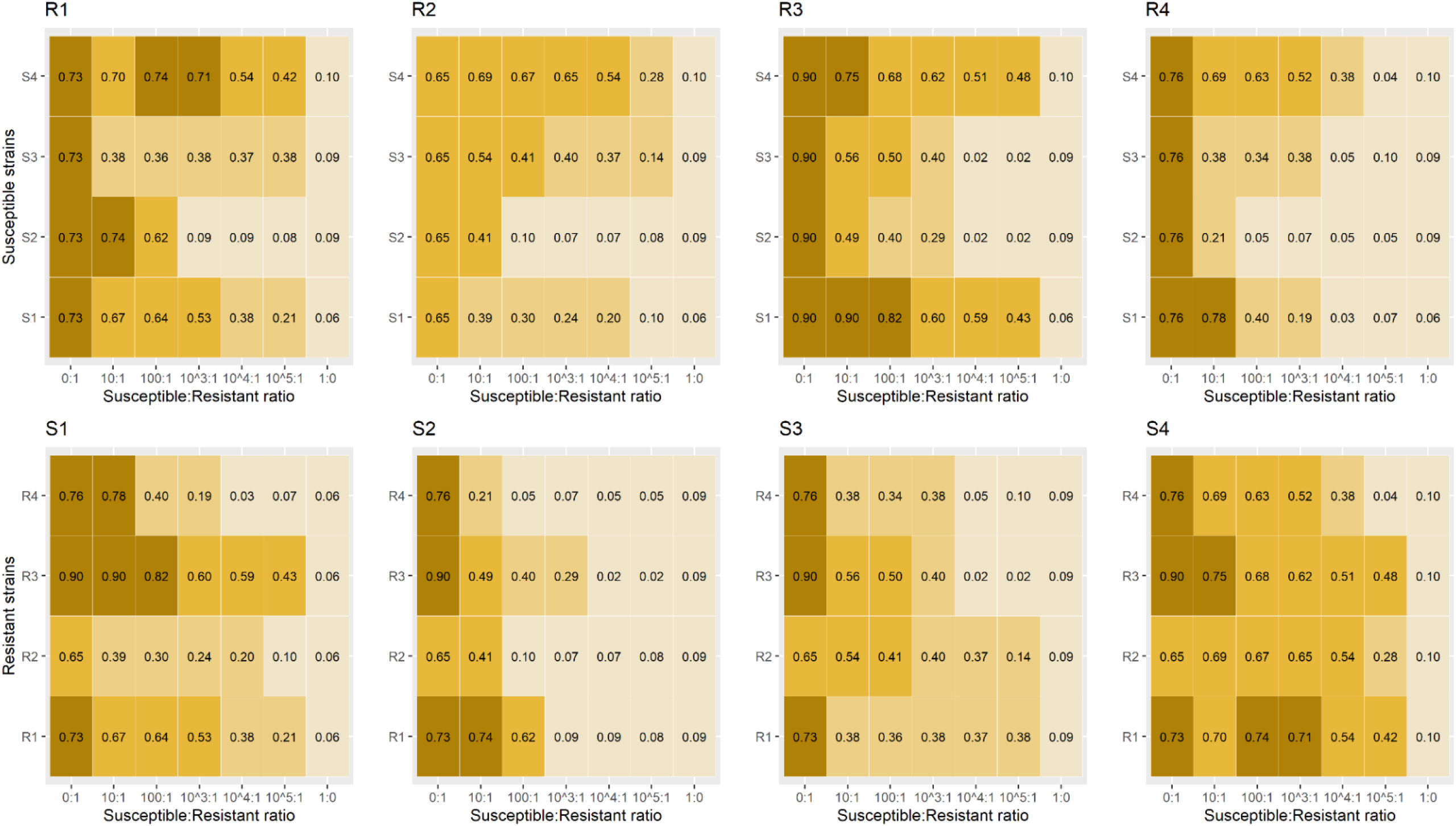
AUC_adj_ of standard suspensions. Positive growth values are represented by increasing strength of color. R= Resistant, S= Susceptible. Average of AUC_adj_ of resistant strain 1, 2, 3 and 4 was 0.40, 0.30, 0.41 and 0.26, respectively. Average AUC_adj_ of susceptible strain 1, 2, 3 and 4 was 0.41, 0.19, 0.31 and 0.54 respectively.

**TABLE 1.** Description of AMU and estimated prevalence of colistin resistance in 36 small-scale chicken flocks stratified by whether farmers administered colistin or not.

|  | Flocks not using<br>colistin (n=14) | Flocks using<br>colistin (n=22) | All flocks<br>(n=36) |
| --- | --- | --- | --- |
| Cycle duration (weeks) (median [1<sup>st</sup> - 3<sup>rd</sup> quartile]) | 19 [17-20] | 20 [17-21] | 19 [17-20] |
| No. chickens (median [1<sup>st</sup> - 3<sup>rd</sup> quartile]) | 249 [194-482] | 208 [128-398] | 231 [189-401] |
| No. ADDs of colistin (per 1,000 chicken-days) (mean ± SD) | | | |
| First period | 0 | 281.7 ± 321.2 | 172.1 ± 285.1 |
| Second period | 0 | 17.4 ± 18.1 | 10.6 ± 16.4 |
| Whole production cycle | 0 | 149.5 ± 261.6 | 91.4 ± 216.4 |
| No. ADDs of non-colistin antimicrobials (per 1,000 chicken-days) (mean ± SD) | | | |
| First period | 345.5 ± 471.5 | 629.3 ± 359.8 | 518.9 ± 424.2 |
| Second period | 29.0 ± 48.6 | 72.5 ± 98.5 | 55.6 ± 84.7 |
| Whole production cycle | 187.2 ± 366.2 | 350.9 ± 383.8 | 287.3 ± 382.9 |
| No. flocks using colistin two weeks prior to |  |  |  |
| Mid-sampling | 0 | 11 | 11 |
| End of sampling | 0 | 1 | 1 |
| AUC<sub>adj</sub> (median, [1<sup>st</sup> - 3<sup>rd</sup> quartile]) |  |  |  |
| Day-olds | 0.07 [0.04-0.42] | 0.06 [0.04-0.52] | 0.07 [0.04-0.65] |
| Mid-production | 0.06 [0.03-0.43] | 0.54 [0.07-0.65] | 0.20 [0.05-0.63] |
| End of production | 0.07 [0.06-0.55] | 0.07 [0.06-0.55] | 0.07 [0.05-0.56] |
| Prevalence of resistance (%) at sample level (95% CI) |  |  |  |
| Day-olds | 42.8 (18.8-70.3) | 31.8 (14.7-54.9) | 36.1 (21.3-53.8) |
| Mid-production | 28.6 (9.5-58.0) | 63.6 (40.8-82.0) | 50.0 (34.5- 65.5) |
| End of production | 28.6 (9.5-58.0) | 31.8 (14.7-54.9) | 30.5 (16.9- 48.3) |
| Estimated prevalence of resistance (%) at colony level (mean ± SD) | | | |
| Day-olds | 28.8 ± 36.0 | 15.7 ± 24.7 | 20.8 ± 29.8 |
| Mid-production | 17.3 ± 28.7 | 27.0 ± 26.4 | 23.3 ± 27.4 |
| End of production | 16.2 ± 24.8 | 12.8 ± 18.1 | 14.1 ± 20.7 |

AUC= area under the growth curve; CI= Confidence interval; SD= standard deviation

In addition to colistin, a total of 28 non-colistin antimicrobials (belonging to 12 classes) were administered to study flocks.. In decreasing order, oxytetracycline, tylosin, neomycin, ampicillin, streptomycin and doxycycline were the antimicrobials most used. The average number ADDs per 1,000 chicken-days of other antimicrobials among flocks using colistin was higher than flocks did not use colistin (350.9 SD ±383.8 vs. 187.2 SD ±366.2, Wilcoxon test, p= 0.004). Among both type of flocks, antimicrobials were administrated more commonly during the first period (average No. ADD per 1,000 chicken-days 629.3 SD ±359.8 and 345.5 SD ±471.5, respectively) compared to the second period of chicken life (average No. ADDs per 1,000 chicken-days 72.5 SD± 98.5 and 29.0 SD ±48.6, respectively) (Table 1).

### Prevalence of colistin resistance at colony level

A total of 909 *E. coli* strains were isolated from 23 selected samples (~40 *E. coli* isolates/ sample) and were tested for their MIC against colistin. Among those, total of 129 strains (14.2%) were resistant to colistin. Of resistant strains, 75.2% strains had a MIC of 4 mg/L, whereas 24.0% had a MIC of 8mg/L. Only 1 isolate (0.8%) displayed a MIC of 16mg/L (Fig. S1). The beta-regression model that relates the AUC_adj_ to percentage of resistant bacteria in samples is shown in Fig. 2. The trend over AUC_adj_ was highly significant (p< 0.001). The equation 100/(1+e^4.8-(7.04*AUCadj)^) associated with this model was applied for estimating the prevalence of colistin resistance at colony level among field samples.

**FIG 2.**
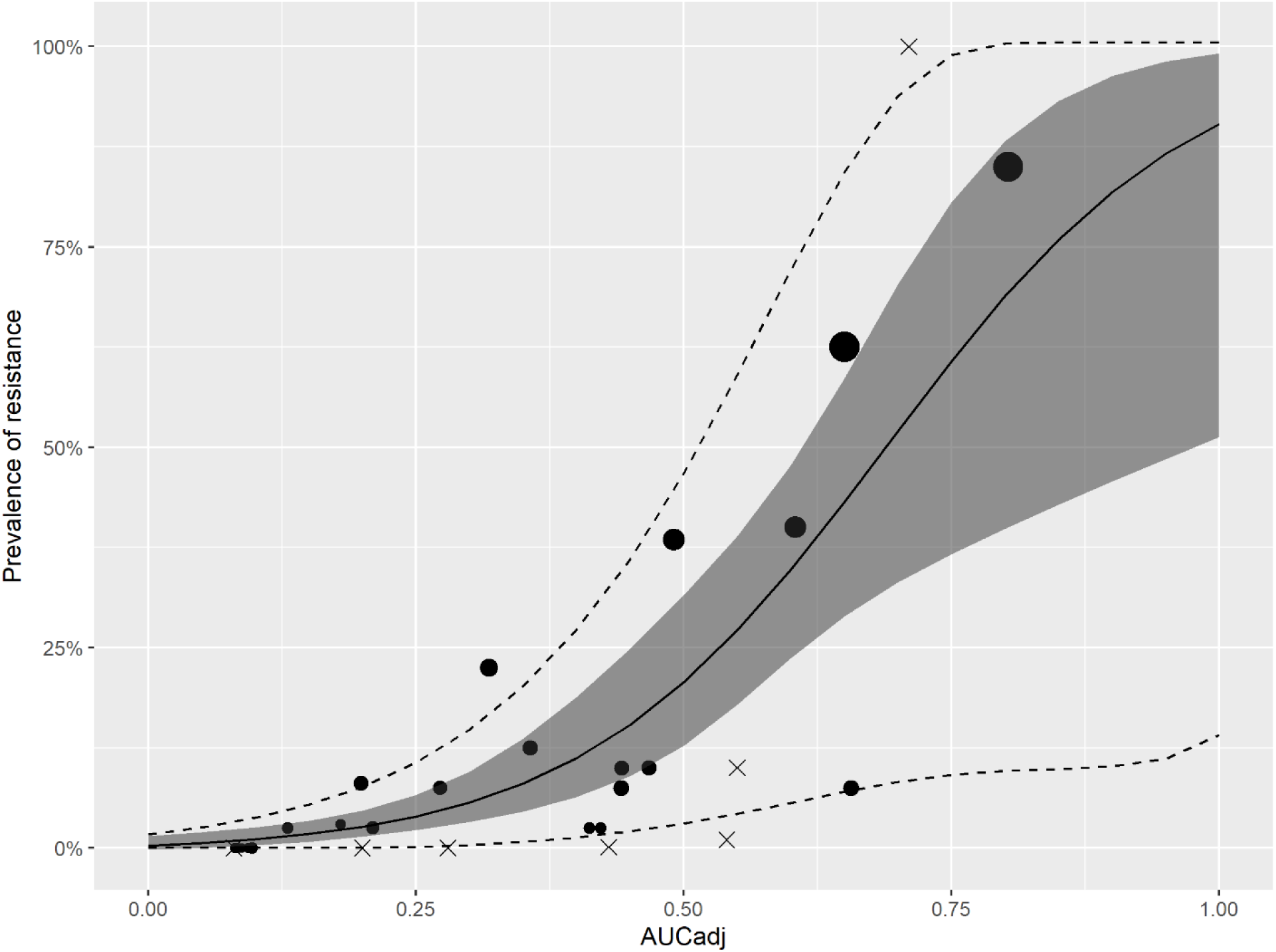
Relationship between AUC_adj_ and prevalence of colistin resistance at colony level. The figure shows the predicted mean value of resistance with pointwise 95% confidence as shaded area. The dotted lines give the 5% and 95% prediction intervals. Circle symbols indicated AUC_adj_ values of mixed *E. coli* in field samples. Size of dot represented the average MIC of each sample. Cross symbols indicated AUC_adj_ values of mixed susceptible and resistant strains.

### Changes of AUC_adj_ over production cycle and prevalence of colistin resistance

Overall, there was no significant change colistin resistance (AUC_adj_) over the production cycle (p= 0.569, Fig. S3). Among flocks not exposed to colistin, the differences AUC_adj_ between sampling points were small. However, among flocks using colistin, the AUC_adj_ values for mid-production samples (0.54 [1^st^ - 3^rd^ quartile, 0.07-0.65]) were higher than those of day-olds (0.06 [1^st^ - 3^rd^ quartile, 0.04-0.52]) (Wilcoxon paired test, p= 0.063) and end of production samples (0.07 [1^st^ - 3^rd^ quartile, 0.06-0.55]) (Wilcoxon paired test, p= 0.046). There was little to no difference in AUC_adj_ values between day-old and end of production samples (Table 1). The prevalence of colistin resistance at sample level was 38.9% (42/108 positive samples). The prevalence of resistance level of day-old, mid-, and end of production samples was 36.1%, 50.0% and 30.5%, respectively (χ^2^ test, p= 0.219). The overall average prevalence of resistance at colony level was 19.4 SD ±26.3%. Among flocks using colistin, the highest level of resistance corresponded to mid-production samples (27.0 SD ± 26.4%), followed by day-old (15.7 SD ± 24.7%) and end production (12.8 SD ±18.1%) (Kruskal Wallis test, p= 0.070). In contrast, among non-using flocks, day-old samples showed higher prevalence of resistance (28.8 SD ±36.0%) compared to mid (17.3 SD ±28.7%) and end production (16.2 SD ±24.8%) (Kruskal Wallis test, p= 0.453). Summary results are presented in Table 1 and individual sample results are given in Table S1.

### Risk factors for colistin resistance

Table 2 shows results for univariable and multivariable analyses. In the multivariable model, use of colistin during the two weeks prior to sampling (OR= 3.67; 95% [Confidence Interval] CI 0.68-19.7) and use of non-colistin antimicrobials (OR= 1.84; 95% CI 0.88-3.85) were associated with colistin resistance at sample level.

**TABLE 2.** Logistic regression models investigating risk factors associated with colistin resistance in chicken flocks at sample level. Models were based on a total of 72 samples (mid and end production); 29 were positive resistance to colistin.

| Variable | Univariable<sup>a</sup> |  |  | Multivariable |  |  |
| --- | --- | --- | --- | --- | --- | --- |
|  | OR | 95% CI | p-value | OR | 95% CI | p-value |
| Age of chicken flock (weeks) | 0.93 | 0.84-1.02 | 0.156 | 1.04 | 0.91-1.18 | 0.605 |
| Use of colistin within last two weeks (Yes/No) | 5.30 | 1.17-24.08 | 0.030 | 3.67 | 0.68-19.70 | 0.128 |
| No. ADDs per 1,000 chicken-day of colistin<sup>b</sup> | 1.66 | 1.00-2.76 | 0.049 | 1.06 | 0.55-2.06 | 0.845 |
| Colistin resistance of day-old chicks (Yes/No) | 1.45 | 0.53-3.97 | 0.461 | 1.61 | 0.54-4.84 | 0.395 |
| No. ADDs per 1,000 chicken-day of non-colistin antimicrobials<sup>b</sup> | 2.10 | 1.18- 3.73 | 0.012 | 1.84 | 0.88-3.85 | 0.102 |

<sup>a</sup>The variable 'Age of chicken flock' was included in all univariable models to calculate estimates for all subsequent variables. <sup>b</sup>logarith transformed after adding 1, ADD= animal daily dose; OR= Odds ratio; CI= Confidence interval.

### Estimation of test costs

The reagent and media costs of broth microdilution and Etest for testing one sample based on the investigation of 10 *E. coli* isolates were ~24.5 and ~63 US dollars (USD), respectively. The cost for testing one sample by the growth-based method (based on 40 isolates) was ~6.5 USD. In addition, broth microdilution involved a higher labour cost (average of ~1 person-day per sample) compared with either the Etest or the growth-based method (~0.5 person-day) (Table S2).

## DISCUSSION

We present here an approach for the phenotypic investigation of colistin resistance in pooled *E. coli* populations from poultry feces using a broth microdilution-based method. Colistin is widely used in poultry and pig production worldwide (4, 15, 16). In the Mekong Delta of Vietnam, colistin is typically administered to chicken flocks in drinking water during the brooding period (1-4 weeks) with a prophylactic purpose (i.e. to prevent disease) (5). Colistin is also included in some pig and poultry commercial feeds as a growth promoter (AGP) (17). However, from 2020 onwards, AGPs are longer be allowed in Vietnam (Law No. 32/2018/QH14), in line with legislative restrictions in Thailand (2015) (18), China (2016) (19) and India (2019) (20). In contrast with the study of human patients, where colistin susceptibility testing is required to inform therapeutic choices (21) our method is aimed at estimating colistin resistance in mixed commensal *E. coli* populations. Through evaluation of the growth curves of standard *E. coli* suspensions from faecal samples, our method enables the detection of colistin resistance in a dichotomous fashion (presence/absence), as well as providing a quantitative assessment of colistin resistance at colony level (prevalence of resistant *E. coli*). The sensitivity of this methodology is, however, limited by the number of colonies harvested per sample (30-50), and may therefore miss colistin resistant strains in situations of very low prevalence. Indeed, statistically, given a sample of 40 colonies, there is a 5% probability of not detecting colistin resistance in any of them when the prevalence of resistant falls below 7.5%. Because of this, the method is more suitable advised for situations of medium to high prevalence of colistin resistance. The sensitivity could however be potentially increased by collecting several samples or increasing the number of *E. coli* colonies used in each suspension. For example, detection of a prevalence of 2% would require the investigation of 150 isolates (~4 samples, each with 30-50 colonies), detection of a prevalence of 1% would require 300 isolates (~8 samples); 0.1% a total of 3,000 isolates (~75 samples).

Although there was a reasonable relationship between prevalence of resistance and AUC_adj_, we observed considerable variation in AUC_adj_ for similar prevalence values both in our laboratory validation as well as on flock samples. This suggests variable growth capacity among resistant strains, which may depend on their relative fitness. In the case of field suspensions containing a diversity of susceptible and resistant strains, it is also likely that the relative composition of strains may result in variable growth among the resistant strains due to the liberation of toxins (i.e. colicins) in the culture media (22). This may also explain the variable limit of detection confirmed in laboratory conditions with different susceptible strains. In general, given identical prevalence of resistant strains, we observed higher AUC_adj_ values for individual susceptible-resistant strain combinations, compared to the mixed of *E. coli* in field samples (Fig 2). It could be probably explained by less competition exerted in mixes containing a single strain, compared with heterogenous mixes containing ~40 different strains. Because of these reasons, prevalence estimates derived from AUC_adj_ should be always interpreted with caution.

We believe that our testing approach is more efficient than isolating and investigating individual colonies, at a relatively lower cost. However, it requires investment on a microplate reader costing between 3,000 and 10,000 USD. The technique presented here could potentially be adapted to the investigation of other types of phenotypic resistance in *E. coli* (i.e. tetracycline, ampicillin, etc.) but it would necessarily require optimizing working concentrations.

At the colony level, we obtained a median prevalence of 19.4% colistin resistance in flocks. These results are comparable with previous studies on chicken *E. coli* isolates in the area (12-22%) (7, 8). Furthermore, the observed ~40% resistance at sample level is consistent with a previous study on chickens in the Mekong Delta of Vietnam, where 5 *E. coli* colonies were investigated from each of 18 faecal samples (8). In such study, a total of 8/18 (44%) samples included at least one resistant strain (NT Nhung, personal communication). A PCR-based study in this region reported that 59.4% chicken samples investigated tested positive for *mcr*-1 gene (7).

We demonstrated a short-term increase in phenotypic colistin resistance following administration of colistin use as well as non-colistin antimicrobials. This contrasts with a study conducted on a broiler flock in France, where administration of colistin failed to induce colistin resistance in Enterobacteriacea (including *E. coli*) (23). However, unlike in Vietnam, colistin use and resistance (including *mcr*-1) is relatively rare in European livestock (10). Overall, we found relatively high levels of colistin resistance (~40%), even in flocks that had not been given colistin (33.3%). There was evidence of colistin resistance in mid-production samples from flocks that had previously tested negative in day-old samples, and had not been administered colistin (3 of 8 flocks) (data not shown). This suggests that colistin resistance may have been generated or introduced to study flocks from other sources, such as contaminated water or feed, or due to contamination with bacteria from other animal species present in these small-scale farms.

Our findings of increased colistin resistance in flocks treated with antimicrobials other than colistin are intriguing. In a previous study on Mekong Delta pig farms, colistin resistance in *E. coli* strains was associated with use of non-colistin antimicrobials such as quinolones and cephalosporins (8). The presence of genes conferring for resistance against several different antimicrobial classes in *mcr*-harboring plasmids may explain these findings, and suggest that the use of non-colistin drugs may also select for colistin resistance (24).

We observed a peak of colistin resistance in mid-production samples among flocks using colistin, and generally levels of resistance decayed subsequently. This is likely to reflect the higher frequency of colistin use during the brooding period. A longitudinal study on travelers colonized by *mcr-1*-carrying bacteria showed that they were able to completely eliminate these bacteria within one month after returning to their home country (25). The reasons for a reduction in resistance over time are known and may be due to a combination of factors leading to plasmid loss and/or fitness costs. However, studies in the laboratory have shown that the presence of plasmid-mediated colistin resistance has been shown to confer no fitness costs to *E. coli* (26). It is worthwhile noting that in our study chicken flocks were of local native breed, and they were typically raised over a 4-5 month period, a period much longer than that required by industrial broilers (typically 1.5 months). This suggests that birds slaughtered earlier may have a higher prevalence of colistin resistance, and this potentially represents an additional risk to the consumer.

In summary, we developed and validated an affordable method that may be effectively used to quantify colistin resistance in commensal *E. coli* in chicken flocks. Our method may also be adapted to benchmark and monitor changes over time in colistin resistance in faecal samples in other complex biological systems such as abattoirs, slaughter-points and sewage, or even in human individuals. Our results indicate a high background of colistin resistance even in flocks not using this antimicrobial. The observed increases after colistin use were short-lived and suggest that in small-scale farming systems reducing colistin resistance may require increasing biosecurity as well as restocking colistin-negative day-old chicks.

## MATERIALS AND METHODS

### Study design

In order to investigate the biological basis and the limit of detection of the proposed method, we used four previously characterized *mcr*-1 colistin resistant *E. coli* strains, two displaying moderate-level (MIC= 8mg/L) and two low-level (MIC= 4mg/L) colistin resistance, alongside four colistin-susceptible strains. We prepared standard bacterial suspensions consisting of a mix of each of the resistant and the susceptible strains at different ratios; these were incubated in medium with and without 2mg/L of colistin. A growth curve from each suspension was obtained by measuring the optical density (OD_600nm_) during incubation. The area under the curve (AUC_adj_) of each colistin-containing standard suspension was adjusted by the AUC values obtained from its equivalent colistin-free suspension. We investigated the relationship between the prevalence of resistance at colony level and the observed AUC_adj_ from the examination of 30-50 individual *E.coli* isolates from each of 23 samples and obtained a model equation. We calculated AUC_adj_ values of suspensions consisting 30-50 *E. coli* colonies harvested from each of 108 pooled faecal samples from 36 small-scale (single-age) chicken flocks raised in Dong Thap province (Mekong Delta, Vietnam) (27). We inferred the prevalence of resistant *E. coli* in flock samples investigated by extrapolation using the model equation. The contribution of colistin use and other antimicrobials administered to flocks on the observed phenotypic colistin resistance was investigated by building logistic regression models with age as primary time variable.

### Culture of standard suspensions and calculation of AUC_adj_

Each of the chosen resistant *E. coli* strains (named R1 to R4, where R1 and R2 had MIC= 4mg/L; R3 and R4 had MIC= 8mg/L) and susceptible (all MIC≤ 1mg/L) strains (S1 to S4) were incubated in cation adjusted Mueller Hinton II Broth II (MHB2, Sigma-Aldrich, USA) at 37^°^C, 200 rpm for 4h (log-phase) and these bacterial inoculum were adjusted to 10^8^ CFU/mL (OD_600nm_= 0.1), and then diluted down with MHB2 to 10^6^ CFU/mL. Each resistant strain was mixed with a susceptible strain (total 16 combinations) at susceptible: resistant ratios ranging from 0:1 (susceptible strain only) to 1:0 (resistant strain only). Intermediate ratios were 10:1, 10^2^:1, 10^3^:1, 10^4^:1, and 10^5^:1. A total of 100*µ*L of each suspension was added into a well of polystyrene microplate (Corning, USA), containing 100*µ*L of colistin solution (final working concentration was 2mg/L). In addition, respective colistin-free (control) suspensions were prepared. Plates were incubated in a microplate reader (SPECTROStar, BMG Labtech, Germany) at 37^°^C for 20h, and the turbidity (OD_600nm_) readings were recorded every hour.. All experiments were conducted in triplicate.

The areas under the curves (AUC) generated over the 20-hour observation period were computed. The AUC value generated from each standard suspension (AUC_[i]_) was related to the AUC generated by its respective colistin-free control (AUC_adj_= AUC_[i]_ /AUC_[0]_).

### Flock sample and AMU data collection

Fresh pooled faecal samples were collected from each flock at three time-points: (1) day-old chicks, (2) mid-production (~2-3 months-old) and (3) end of production (~4-6 months-old). Day-old faecal (i.e. meconium) samples were collected from the crates at the time when chicks were delivered to the farms. For mid- and end-production sampling, sterile paper liners were placed near drinkers and feeders in the chicken house/pen to collect deposited droppings. After a minimum of 10 droppings had been deposited, liners were swabbed using sterile gauzes. Each of collected gauze was placed in a universal jar and mixed vigorously with 50mL saline buffer. One ml of the resulting eluate was stored at −20^°^C with glycerol. Data on AMU had been collected using purposefully designed diaries where farmers were asked to note down all antimicrobials used. Farmers were instructed to keep all packages of antimicrobials used on their flocks (5). Sample and data collection were conducted between October 2016 and October 2018.

### Testing of pooled faecal samples

Eluates from pooled faecal samples were plated onto ECC agar (CHROMagar, France) and incubated at 37^°^C for 20h. A total of 30-50 *E. coli* (blue) colonies from each agar sample were picked, pooled and incubated in CAMHB to log-phase. The resulting bacterial suspensions were investigated as described above.

### Prevalence of colistin resistance at colony level

We selected a number of positive samples with variable levels of AUC_adj_. From each sample, 40 *E. coli* were isolated and tested individually for the MIC by standard broth micro-dilution method. Pool of 40 isolated *E. coli* for each sample was also subjected for the growth curve measurement as described previously.

### Data analyses and cost estimation

In order to relate the AUC_adj_ value to the measured prevalence of resistance among selected samples, we fitted a beta-regression model using the ‘*betareg*’ package in R (28). Both the trend and the dispersion were allowed to vary over AUC_adj_ in a linear way.

AMU in flocks was quantified for the two periods defined by the sampling schedule: (1) between restocking and mid-production, and (2) between mid- and end of production. Weekly estimates of colistin use were expressed as the number of ADDs (number of Animal Daily Doses administered per 1,000 chicken days) calculated for each of the two periods (5). Risk factors associated with colistin resistance at mid- and end of production were investigated by logistic regression. The outcome was colistin resistance (Yes/No) at sample level. The variables investigated were: (1) Age of chicken flock (weeks); (2) Use of colistin within two weeks prior to sampling (Yes/No); (3) Number of ADDs per 1,000 chicken-days of colistin in each period; (4) Colistin resistance of day-old chicks (Yes/No); and (5) Number of ADDs per 1,000 chicken-days of non-colistin antimicrobials used in each period. The variable Age of chicken flock was included in all univariable models because it is the principal time variable. Since we had two measurements per flock (mid and end cycle samples), we used generalized estimation equations with an exchangeable correlation structure to estimate the parameters using the ‘*geepack*’ R package (29, 30).

The change in AUC_adj_ over age of chicken was modeled using a random effects linear regression. In order to allow for a nonlinear trend, we used a natural spline for the fixed effect term (knots at 0, 8, 12 and 20 weeks). We allowed for a random intercept and linear trend by age.

The overall costs (per sample) of the method described above were calculated based on expenses on medium, reagents and consumables (excluding staff time, which was estimated separately). The estimated costs were compared with those incurred in testing one sample by broth microdilution and Etest in Vietnam as of January 2020. Our calculations were based on the investigation of 40 *E. coli* isolates per sample using the growth-based method, compared with 10 isolates each by broth microdilution and by Etest.

## SUPPLEMENTAL MATERIAL

Table S1 Estimated percentage of resistant *E. coli* from 108 samples

Table S2 Estimated costs (in US dollar) of testing 1 sample to determine colistin phenotypic resistance of *E. coli*

FIG S1 Usage of colistin among study flocks by week.

FIG S2 Changes in AUC_adj_ by age (weeks) of chicken at the time of sampling.

## ACKNOWLEDGEMENT

We would like to thank staff at the Sub-Department of Animal Health and Production (Dong Thap) for their supports in sample and data collection. This work was funded by the Wellcome Trust through an Intermediate Clinical Fellowship awarded to Juan Carrique-Mas (Grant No. 110085/Z/15/Z).

We declare that we have no competing interests.

N.T.N and JC-M conceived the idea; J.C, G.T and S.B advised on the study design; N.V.C and B.T.K coordinated field sampling and data collection; N.T.N, N.T.P.Y. and N.V.K.T performed laboratory experiments. N.T.N, R.B.G and J.C-M conducted data analyses and produced first draft. All authors commented on subsequent versions.

